# Detection of genetically modified organisms using highly multiplexed amplicon sequencing

**DOI:** 10.1101/2023.12.22.573157

**Authors:** C. Sarai Reyes-Avila, Dominique Waldvogel, Nicolas Pradervand, Sylvain Aubry, Daniel Croll

**Affiliations:** Laboratory of Evolutionary Genetics, Institute of Biology, University of Neuchâtel, CH-2000 Neuchâtel, Switzerland; Department of Evolutionary Biology and Environmental Studies, University of Zürich, Zürich, Switzerland; Agroscope. Posieux, Switzerland; Department of Plant and Microbial Biology, University of Zürich, Zürich, Switzerland; Federal Office for Agriculture, Bern, Switzerland

**Keywords:** genetically modified organisms, crops, multiplex PCR, NGS, amplicon-sequencing

## Abstract

The circulation of products based on genetically modified (GM) organisms is highly regulated by some governments having implemented strict rules on the breeding, planting, marketing, labelling, and trading of such products. To ensure compliance, accurate detection methods for GM events are necessary, along with assurance that GM material falls within relevant threshold levels. The increasing complexity and potential of undocumented GM are a growing challenge for genetic screening. In this study, we developed and assessed a highly multiplexed amplicon sequencing assay for the detection of GM events based on a microfluidics platform and next-generation sequencing (NGS). To probe GM events comprehensively, we designed a total of 230 new amplicons to cover flanking, promoter, junction and coding sequences of GM sequences. In addition, we designed and implemented parallel amplification of ribosomal and chloroplast markers to define crop species identity from potentially mixed samples. Using reference GM material of 11 crop species and multiple amplicons, we successfully detected the presence of 10 known modifications per GM event. We also find that reported flanking sequences of GM events may not be all useful for diagnostic. We assessed the assay’s potential to detect GM events in mixed samples as well as in highly diluted DNA. Finally, we performed a prospective search of potentially undocumented GM events in plant material. Our microfluidics-based amplicon GM detection approach fills important gaps in detecting potentially undocumented and complex GM events by recovering a wide range of specific amplicon sequences for evaluation. Integrating highly parallel amplicon assays in GM screening efforts should be an effective complement to aid post-market monitoring and regulatory compliance efforts.

## INTRODUCTION

Genetically modified (GM) crops contain genes that have been artificially introduced conferring beneficial traits such as herbicide tolerance, and drought or pest resistance (Alasaad, Alzubi, and Kader 2016). The production of GM organisms has been rising and becoming more widely available commercially. Commercial applications of GM in crops have led to hundreds of variants of genetic modifications in dozens of crop species but mostly in the main cash crops (maize, soybean, canola and cotton. Since the‘90s, the European Union (EU) implemented risk-based legislation governing the planting, marketing, labelling and trade of GMOs in Europe (Serageldin 1999). To enforce legislation, it is essential to develop accurate methods for the detection of GM and monitoring of threshold levels.

Any material deriving from GM crops might be identified by testing for the presence of introduced DNA. GM events or constructs that are often found in commercialized GM crops usually consist of several elements (promoters, terminators, genes, antibiotic resistance cassettes) that help to screen for their presence. There is a broad range of methods for GM DNA detection, and quantitative polymerase chain reaction (qPCR) approaches remain the most common (Aubry et al. 2021). qPCRs allow the detection, identification, and quantification of genetic modifications, based on PCR amplification recorded in real-time. As an example, a semiautomated TaqMan PCR screening of GMO-labelled samples was recently developed (Scholtens et al. 2017). Scholtens *et al*. semiautomated screening based on 32 primer and probe sequences derived from methods used in routine GMO feed sample screenings. Scholtens *et al*. verified the 32 primers in 59 different GMO reference materials. When a GMO element cannot be explained by any sample labelling, this is indicative of an unknown (*i.e.* unauthorised) GMO. To facilitate the detection of several DNA targets in a single reaction, multiplex PCR-based methods were developed. qPCR-based multiplexing is constrained through costs and technical challenges to retain high reproducibility across laboratories. Official control laboratories are required certification to ensure high reproducibility standards. In addition, strategies have been developed to increase assay throughput without compromising sensitivity or specificity. Next-generation sequencing (NGS) has been used for GM detection, offering new avenues for improving throughput, sensitivity, and scalability. NGS-based approaches allow simultaneously obtaining multiple sequences and may facilitate the unambiguous detection of GMOs (Arulandhu et al. 2018). Arulandhu *et al*. tested five feed samples known to contain GMOs and screened 96 GMO-specific targets including endogenous, elements, constructs and different events. Based on a related approach, Scholtens *et al*. targeted sequences from certified reference materials. Targeted sequencing was mainly applied to samples with multiple GM sequences of interest (Jagadeesan et al. 2019). DNA libraries were obtained from PCR-based amplification including several amplicons that can be sequenced using NGS technology. Standardized bioinformatics pipelines were applied to raw reads for filtering and assembly into contigs, which can be mapped to a reference sequence database (Willems et al. 2016). Establishing reference databases remains challenging as public disclosure of modified sequences can be incomplete (Moreira, Carneiro, and Pereira 2017b).

Here, we developed a highly multipled amplicon sequencing assay for the detection of GM events in crops. The is assays was conceived by integrating a large set of available GM sequences from public databases to design new, robust amplicons to amplify sequences linked to GM events in segments. Barcoding primers were designed to detect the species identity present in the material. Amplifications were conducted in parallel using a microfluidics-based multiplex PCR and amplicons were sequenced as a pool using Illumina NGS. Filtered reads of positive and negative control samples were mapped to a compiled GM amplicon database. The performance of the assay, in particular specificity and sensitivity, in diluted and mixed samples were evaluated.

## MATERIALS AND METHODS

### Collection of samples

Genotyping was performed on 92 plant samples consisting primarily of GM-certified reference materials (Table S1). Samples covered eleven of the most widely cultivated plant species worldwide, such as maize *(Zea mays)*, soybean *(Glycine max)*, canola *(Brassica napus)*, cotton *(Gossypium sp)*, alfalfa *(Medicago sativa)*, potato *(Solanum tuberosum)*, beetroot *(Beta vulgaris)*, creeping bentgrass *(Agrostis stolonifera)*, linseed *(Linum usitatissimum)*, wheat *(Triticum sp)* and rice *(Oryza sp)*. The samples also included non-GMO crops used as negative controls. To assess the sensitivity of the assay, different concentrations of the same GM crops were used, typically ranging from 0.98 % to 100 % (ratio of the GM plant species in the total sample, expressed in mass/mass or copies/copies of haploid plant genomes). Detailed information about each sample and GMO events are provided in Table S1.

### DNA extraction

DNA extractions were carried out using the NucleoSpin Plant II kit (Macherey-Nagel, GmbH, Germany) following the manufacturer’s protocol. DNA concentrations of all samples were assessed using a NanoDrop One spectrophotometer and a Qubit (Thermo Scientific). Initial DNA concentration are reported in Table S1. We chose to standardise the DNA concentration of all our samples by diluting them in water to 50 ng μl^−1^. To explore the effects of low DNA input, we diluted one of the GMO maize samples in a dilution series starting from 50 ng μl^−1^ down to 5 ng μl^−1^ (10-fold), 0.5 ng μl^−1^ (100-fold) and 0.05 ng μl^−1^ (1000-fold). For downstream applications, we performed two additional replicates for 41 samples, including replicates of the dilution series (Table S1).

### Recovery of target sequences and barcoding plant species

We recovered GM sequences as target sequences for amplicon design from two GMO databases: EUginius (EUropean GMO INItiative for a Unified Database System) and portugene (Moreira, Carneiro, and Pereira 2017a). Additionally, we added sequences encoding the dihydroflavonol 4-reductase gene from an unauthorized GM *Petunia* sequence (Fraiture et al. 2019). To avoid redundancy between the target sequences, we clustered nearly identical sequences and produced multiple alignments using Clustal Omega-v1.2.3 (Sievers and Higgins 2014). Aligned sequences were used to produce a consensus, retaining ambiguous bases and yielding a total of 115 unique sequences targeting specific GM events (Supplementary File S2). To identify the plant species of tested samples, we added two sets of primers (Kress 2017) widely used as plant DNA barcoding loci for taxonomic identification. RbcL (the chloroplast-encoded large subunit of the Ribulose-1,5-bisphosphate carboxylase-oxygenase) and ITS (internal transcribed spacer) from the ribosomal DNA locus.

### Amplicon design

For the 115 unique target sequences, we segmented sequences exceeding 300 bp into multiple regions for individual amplicon design to improve amplification yields and coverage of the GM events. After the segmentation, we obtained 230 candidate loci for primer design according to Fluidigm Inc. recommendations. The targeted amplicon length ranged from 62-240 bp reflecting constraints in conserved sequences and base composition. We obtained a total of 230 pairs of primers corresponding to the GM target sequences (Supplementary File S1). In the case of the barcodes that amplify rbcL and ITS genes, we compiled a set of 27 primers representing various amplicon designs covering the same loci corresponding to rbcL and ITS (Supplementary File S3).

### DNA sequencing library preparation

Libraries were prepared following the manufacturer’s protocol PN 101-0414 G1 for the Juno LP 192.24 integrated fluidic circuits plate (IFC; Fluidigm Corporation, San Francisco, CA, United States). After loading all reagents on the IFC, target amplicons were generated for each sample through PCR amplification on a specialized thermocycler (Juno system; Fluidigm). A total of 257 primers subdivided into 10X assay pools were combined in the IFC with an inlet containing 2 µl of sample pre-mix, 2 µl genomic DNA (50ng/µl) and 1 µl barcode primer mix consisting of a DNA sample and an individual barcode. After amplification, samples were pooled in a single tube and purified. The first clean-up is double-sided (0.4X/0.9X Double-Sided SPRI), with first (0.4X) removing fragments that are bigger than the targets, then (0.9X) binding and selecting targets by washing off the smaller fragments. The second and third clean-ups were to remove excess primers (0.8X SPRI). Finally, sequencing adapters were added by PCR to the purified library followed by a final round of purification according to the manufacturer’s protocol. The quantity and quality of the library were assessed using a Qubit fluorometer assay (ThermoFisher) and a 4200 TapeStation electrophoresis instrument (Agilent). The final library was sequenced on a single lane of a NextSeq 500 system (Illumina) in mid-output mode adding ∼30% PhiX to reduce issues due to low sequence complexity.

### Recovering of plant barcode sequences and GM targeted sequences by mapping

We obtained the rbcL sequence of maize from NCBI (NC_001666.2) and the ITS sequence from potato (CP046695.1). Matching rbcL and ITS sequences were then retrieved using BLAST (Altschul et al. 1990) to complete a library of barcoding sequences for all eleven included plant species. The crop barcode sequences were added to the GM reference sequences. We demultiplexed raw read data using bcl2fastq v-2.19.0.316 and used trimmomatic v-0.36 (Bolger, Lohse, and Usadel 2014) for quality trimming. Forward and reverse reads were merged using flash v-1.2.11 (Magoč and Salzberg 2011). We aligned merged reads to the reference sequences using bowtie2 v-2.3.5 using the following settings: --very-sensitive-local --phred33 (Langmead et al. 2019). Based on the aligned reads, we calculated the depth per position ignoring locations outside of designed amplicons. We accessed all nucleotide sequences for rbcL and ITS available on NCBI for each of the eleven plant species generating eleven fasta files for rbcL and eleven fasta files for ITS. The fasta files were used as a reference to align the merged reads by bowtie2 v-2.3.5 using the following settings: --very-sensitive-local --phred33 (Langmead et al. 2019). From the aligned reads, we calculated the depth per position using samtools v-1.19. and we compared the depth to the previous depth of the rbcL sequence from maize (NC_001666.2) and the ITS sequence from potato (CP046695.1) with the retrieved BLAST sequences of the eleven crops. NC_001666.2 and CP046695.1 reference sequences were compared based on mapped reads per sample.

## RESULTS

### Design and amplicons quality from the amplicon-sequencing assay

In order to test the feasibility and assess the performances of the amplicon-sequencing assay to detect GM material in food, feed or seed matrices, we gathered a total of 115 consensus GM sequences from various sources: EUginius (EUropean GMO INItiative for a Unified Database System, and portugene (Moreira, Carneiro, and Pereira 2017a) databases (Figure 1A). In addition, we manually added three sequences from a gene encoding the dihydroflavonol 4-reductase from an unauthorized GM *Petunia* recently detected in the (Fraiture et al. 2019). For the design of amplicons, we retrieved the GM consensus sequences if multiple redundant sequences were found in databases. Furthermore, sequences longer than 300 bp were fragmented to design multiple similarly spaced amplicons (Figure 1C). Based on this GM sequence set, 230 primer pairs for GM sequences that correspond to 82 unique GM sequences were designed (Figure 1C) and 15 rbcL + 12 ITS primers were added for plant species identification (Figure 1B). Following the manufacturer’s instructions, we aimed for an amplicon length of ∼200bp and we obtained a maximum length of 240 bp for 14 GM locus and the smallest amplicons with a length in the range of 69-199 bp for 21 GM loci (Figure 2A).

**Figure 1.**
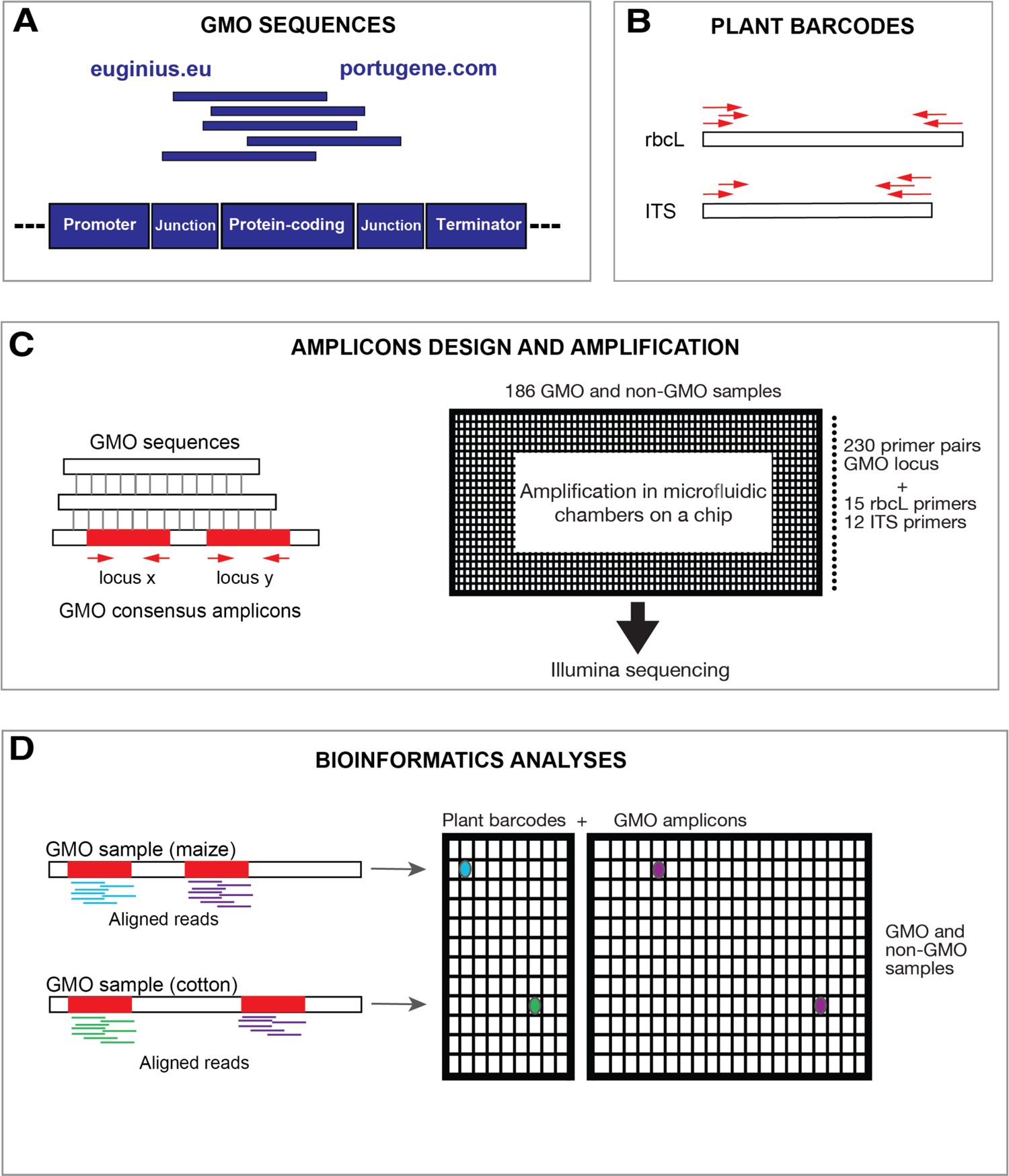
Design and workflow of the targeted amplicon sequencing loci. (a) Recovery of GM sequence events from euginius and portugene databases, and the general structure of a GM event, consisting typically of a promotor, junctions, protein-coding and terminator. Alignment of similar GMO sequences to obtain consensus, fragmented by locus to obtain amplicons around 300bp,) and the microfluidics-based Juno system approaches. Bioinformatics based on alignments of every sample map to plant barcode locus and GMO amplicons. Abbreviations: RBCL, ITS, GMO

**Figure 2.**
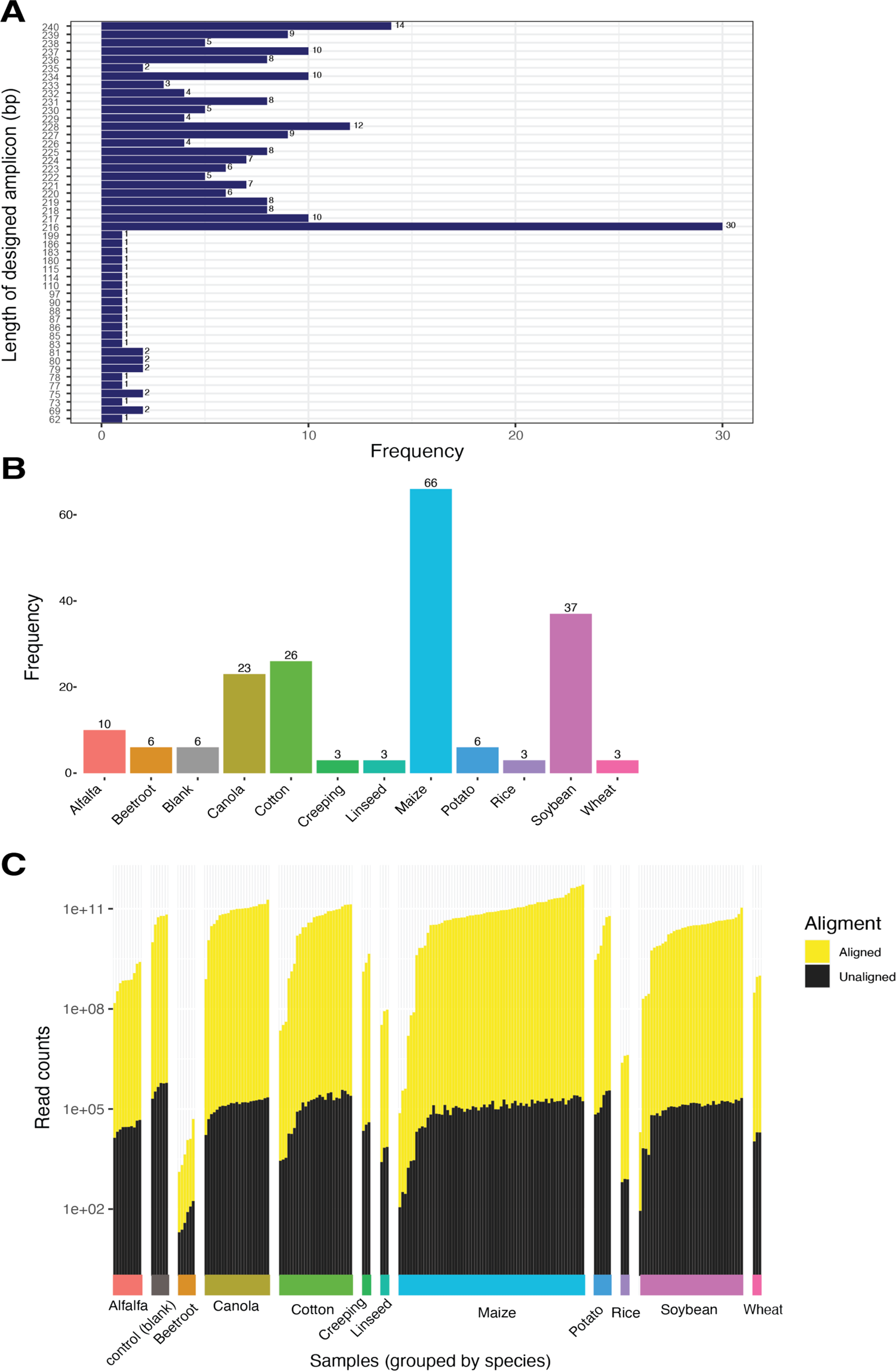
Overview of the designed amplicons, samples used for testing and amplification. (a) Length of the 230 designed GM amplicons. (b) Number of samples per plant species, giving a total of 192 samples that include 92 different individuals GMO and controls, dilutions and replicates. (c) Mapped and unmapped reads per individual sample according to species identity.

The amplicon sequencing assay was performed using a microfluidics platform (Figure 1C). In a single flowcell run, we analysed a total of 186 samples including replicates, representing 92 distinct samples of GMO and non-GMO control material. The species covered were alfalfa (n=10) *(Medicago sativa)*, beetroot (n=6) *(Beta vulgaris)*, canola (n=23) *(Brassica napus)*, cotton (n=26) *(Gossypium sp)*, creeping beetroot (n=3) (*Agrostis stolonifera*), linseed (n=3) *(Linum usitatissimum)*, maize (n=66) *(Zea mays)*, potato (n=6) *(Solanum tuberosum)*, rice (n=3) *(Oryza sativa)*, soybean (n=37) *(Glycine max)* and wheat (n=3) *(Triticum sp)*, additionally blank (n=6) (Figure 2B).

Illumina NextSeq 550 sequencing generated a total of 80,478,507 reads after trimming and merging read pairs per sample and locus. Furthermore, for every sample, the reads were aligned to the 115 target sequences and plant barcoding sequences rbcL and ITS sequences of the 11 species. One sample of maize obtained a maximum of 3,122,899 aligned reads (Figure 2C). The minimum number was 229 aligned reads which corresponds to a sample of soybean (Figure 2C). The other three minima of aligned reads ranged from 663 to 1,418 and correspond to 3 replicates of DNA dilution (1000x) of a maize sample (Figure 2B).

### Identification of plant species by amplification of plant barcoding loci

An important step for the identification of GMO crops is to determine species identities. We incorporated in our targeting sequencing assay two sets of primers (Kress 2017) previously recognized as plant DNA barcodes that amplify the rbcL and ITS loci. Given that our 15 rbcL + 12 ITS primers are amplifying in different regions of rbcL and ITS (Figure 3A), we decided to work with the maximum number of mapped reads per barcode, which were correlated to the mean (*r* = 0.897***, Pearson correlation coefficient), median (*r* = 0.848***) and mode (*r* = 0.848***). To evaluate the success of rbcL and ITS amplification, we extracted the maximum number of mapped reads for the 186 samples. The rbcL maximum count was significantly positively correlated to ITS maximum counts for alfalfa, canola, cotton, potato and soybean in the range of 0.983*** (*p*-value <0.001) to 0.748*** (Figure 3B). In the cases of beetroot, creeping beetroot, rice and wheat, the correlation was not significant (*p*-values > 0.05), given that we have a low number of samples (*n* = 3-6; Figure 3B). Maize showed also a positive correlation with 0.503*** (Figure 3B). We also compared the maximum read count at barcodes against the total aligned counts to ensure that the general quality of the samples has been captured by the plant barcodes. For ITS, 10 plant species showed a significant and positive correlation from 0.88*** to 1***, with the only exception of beetroot lacking a strong correlation (0.25*). RbcL varies more between species with a range of −0.96 p=0.19 to 0.97***.

**Figure 3.**
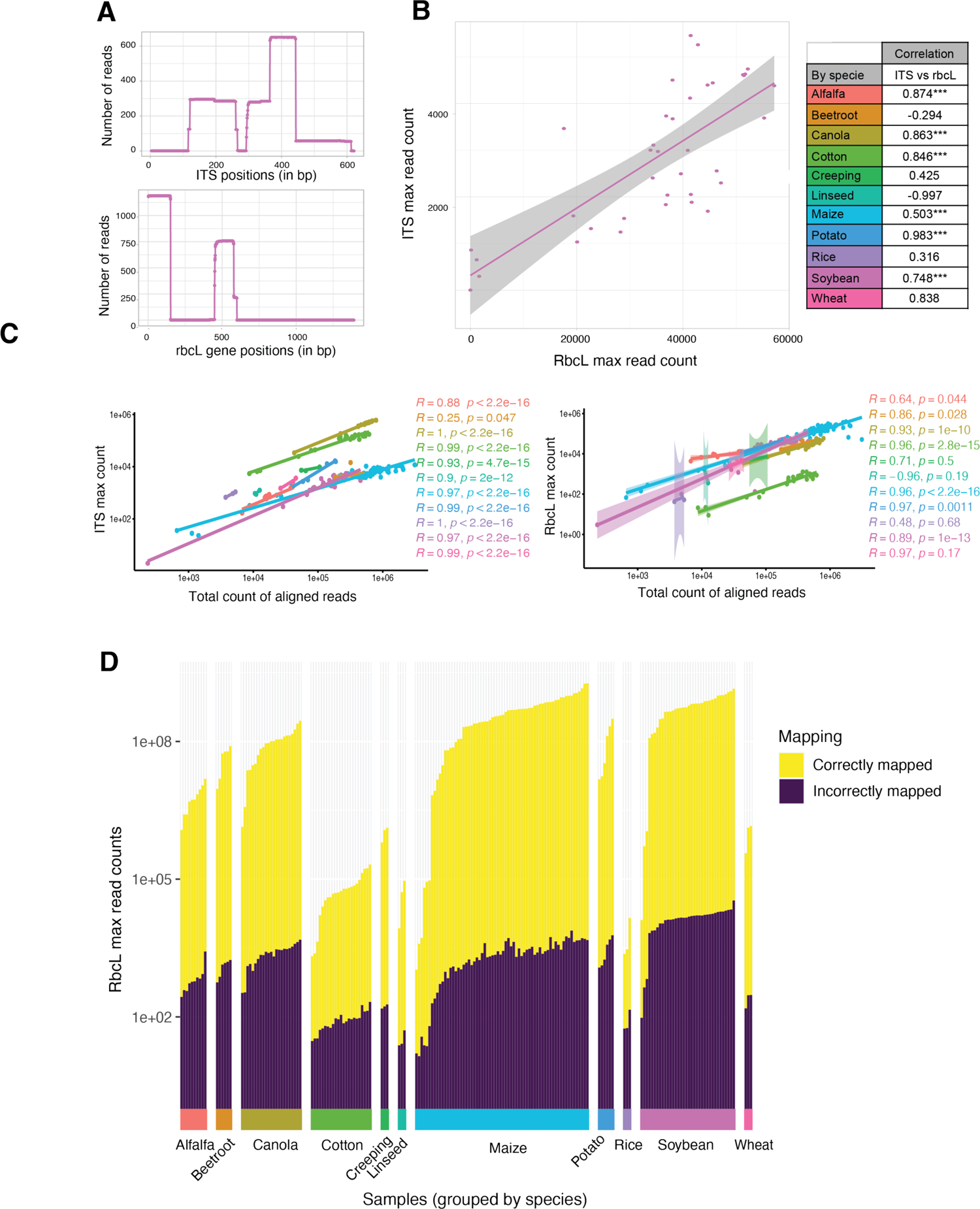
Quantification of read mapping on reference sequences of the ITS and rbcL loci following the amplification using plant barcoding primers. (a) Depth of mapped reads at the ITS and rbcL loci for a soybean sample. (b) Correlations of the maximum read count for ITS compared to rbcL for all soybean samples including a table of correlations for other plant species. (c) Correlation of the total count of aligned reads compared against the maximum count of ITS reads. Correlation of total aligned read count compared against the maximum count of rbcL reads with *r* values and *p-*values. (d) Maximum counts of mapped reads for the rbcL locus distinguishing reads mapped to the correct species.

Every sample was mapped to the 11 crop species sequences of rbcL and ITS. We assigned species identities according to the highest read depth among rbcL and ITS reference sequences. As the species was known for all samples, we assessed the power of the assay to recover the true species. Only using, the ITS primers we achieved 91 out of 186 correct predictions of the species identity corresponding to an accuracy of 48.9%. For the rbcL primers, we achieved 182 out of 186 correct predictions of the species identity corresponding to an accuracy of 97.8%. Hence, rbcL primers predicted more accurately the species and we used this barcoding locus for most species (Figure 3C). The species that performed best was maize showing the highest correctly mapped reads ratio for 13 samples (79-172), meaning that the primers used to amplify rbcL and the rbcL sequence used to align are providing the best correct mapping. The worst correctly mapped reads ratios correspond to the three rice samples: 0.912-0.686.

### Detection of GM events based on amplicon-sequencing assay

We analyzed 98 individual samples including 11 non-GMO controls. The samples were known to carry at least 10 sequence fragments from a specific GM event with 9 samples originating from maize and one from cotton (Supplementary Table S2). We analyzed the 10 GMO samples and compared these against the non-GMO controls. To account for variation in total reads among samples and the uneven presence of GM sequences, we normalized read counts using normalized mean depth by the maximum read depth at the barcoding locus. In the case of maize, we used rbcL because the read depth was higher in comparison to ITS (Figure 3C). For cotton, we used ITS because the read depth was higher for ITS (Figure 3C). From the 10 analyzed GM events, six events (MON89034, BT11, DAS59122, MIR162, MON88017 and MON87427) amplified well in the GMO sample and showed no meaningful amplification in the non-GMO control (Figure 4A-F). For example, MON 89034 amplified well for 3’MON89034 (normalized count 0.094), 5’MON89034 (normalized count 0.004), hsp70-locus56 (normalized count 0.747), hsp70-locus307 (normalized count 0.285), LTa.lhcbl (normalized count 0.002), tahsp17 (normalized count 0.331) and CTP2 (normalized count 0.405). For the three other GM events (MON810, SYN3272 and MON87460), the GMO maize also amplified more GM sequences compared to the non-GMO control (Figures 4G-I). For example, the sample MON810 where the non-GMO control is amplified in 5’ MON810-locus 59 (normalized count 0.383) vs the GM sample (normalized count: 0.526) (Figure 4G). The last sample corresponds to MON15985 where the sequence labelled “Cotton_MON15985” amplified more in the GMO cotton (0.148) compared to non-GMO cotton (0.046 normalized counts; Figure 4J). The most discriminant amplicon for detecting the MON15985 event was the Oriv amplicon.

**Figure 4.**
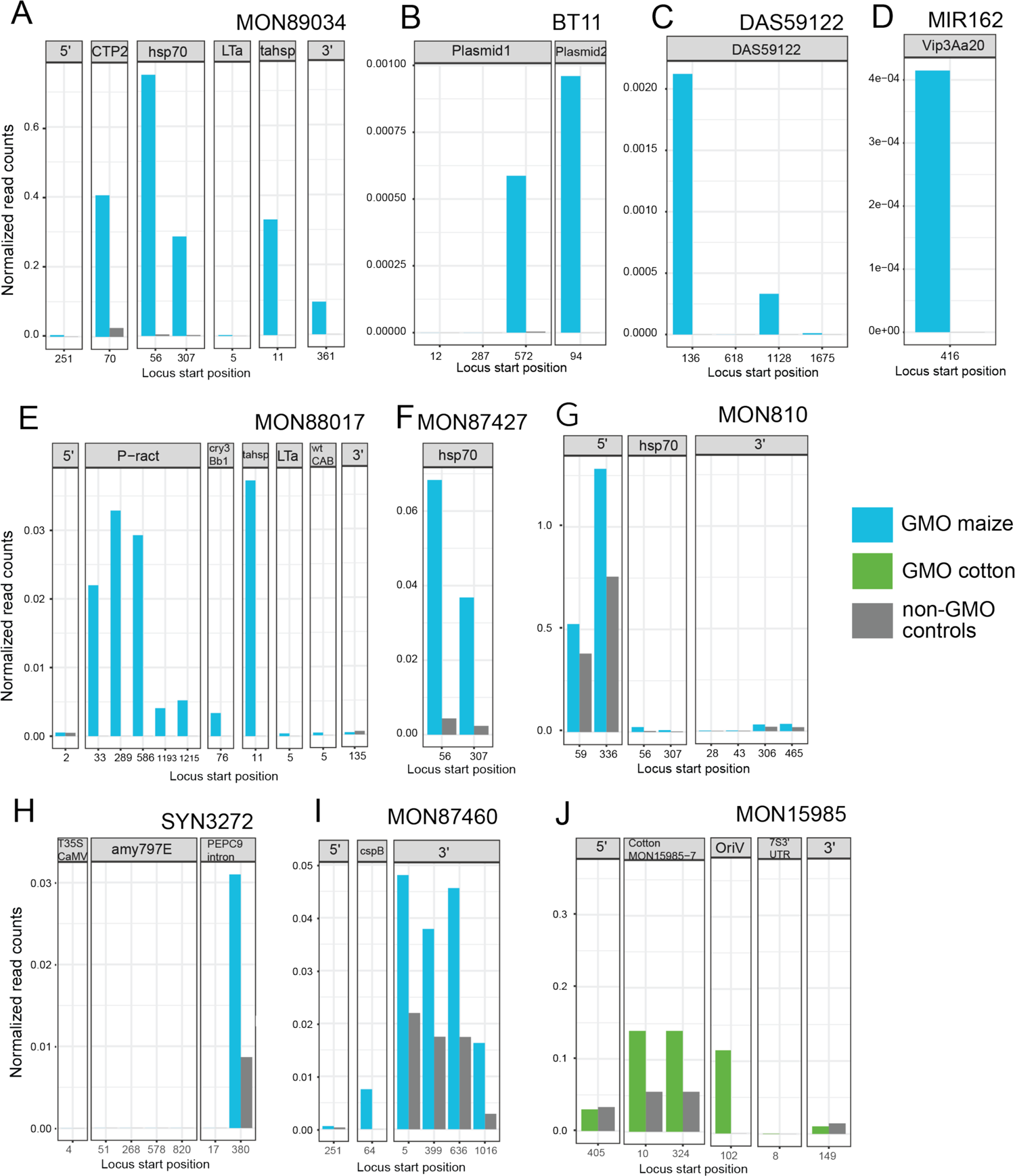
Assessment of known GMO events by amplicon sequencing and contrast to control samples. The following GM events were covered for multiple segments (with base pair positions indicated for multiple amplicons covering the same segment of an event. Events (a) MON89034, (b) BT11, (c) DAS59122, (d) MON88017, (e) MON88017, (f) MON87427, (g) MON810, (h) SYN3272, (i) MON87460 and (j) MON15985.

### Specificity and sensitivity of the detection of the amplicon sequencing assay

For monitoring purposes, mixed GMO samples need to be screened. To assess the power of our targeted sequencing assay, we tested three GMO samples diluted in control DNA at various ratios. For the GMO SYN3272 at the locus amy797E, the GMO (constituting 9.8%) amplified in the range of 0.01-0.002 normalized read counts contrasted with no recovered reads for the GMO mixed in at 0.98% (Figure 5B). The sample DAS59122 amplified in both the 1% mixture in the range of 1.15 e-10 – 0.0021 and the 10% mixture in the range of 0.00018 – 0.00796 (Figure 5C). Unexpectedly, MON810 showed more amplification for GMO 1% compared to GMO 10% (Figure 5A). For example, 5’ MON810 locus 336 amplified for GMO 10% 1.2830 compared to 1.553 normalized read counts amplified for GMO 1%. The non-GMO control sample showed however 0.7572 normalized read counts. Next, we assessed the sensitivity of the amplicon assay to detect DNA diluted 10x, 100x, and 1000x. We tested the sample MON88017, however, the normalized counts mismatched the expected trend from the dilutions. The most likely explanation for this variability is the overall low number of reads likely inducing noise in read counts (Figure 5D). Using the mean depth of mapped reads (before normalization), we recovered the expected change in read depth along the dilution series (Figure 5E).

**Figure 5.**
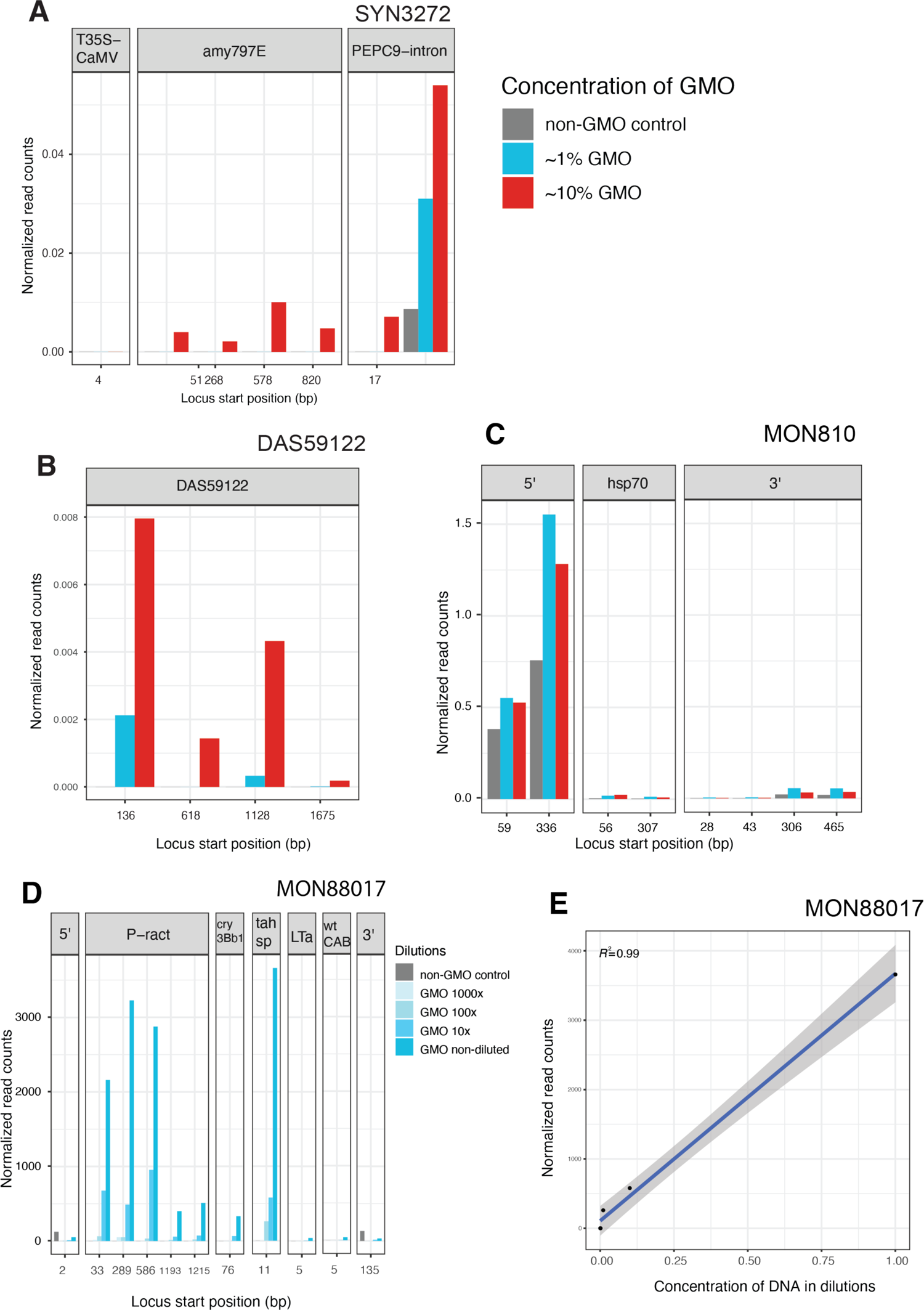
Assessment of different concentrations of GM and dilutions of sample DNA. (a) GM event MON810 spanning three assessed regions (3’, 5’ and hsp70). (b) GM event called SYN3272, composed of three recovered regions (amy797E, PEPC9-intron, and T35S-CaMV) in different locus positions where the reads were aligned and we normalized the count in bp, for the 0.98% concentration of GMO in blue, the 9.8% concentration of GMO maize in red and the non-GMO maize in grey. (c) GM event called DAS59122 amplified at four different loci.

### Additional amplicons for exploratory detection of GM events

We recovered 73 target sequences from EUgenius, portugene and Fraiture *et al*. 2019 and amplified these in using the custom amplicon sequencing assay. We assessed amplification of the 10 GM sequences described above (Figure 6). MON15985 cotton is amplifying the sequence Cotton_MON15985 as expected and also OriV (Figure 4J). Non-GMO controls showed amplification of multiple 3’ and 5’ GM flanking sequences as expected from the above findings. Three additional sequences including “FB707511.1 cry1A.105”, “DL476427.1 CORN EVENT” and “aadA” showed amplification in the MON15985 GMO. In regard to the GMO maize samples, MON 89034 amplified the sequences 3’MON89034, hsp70, tahsp and CTP2 (Figure 4A). Beyond amplified flanking sequences, three sequences amplified in the GMO including “FB707511.1 cry1A.105” and “DL476427.1 CORN EVENT”, and “dihydroflavonol4-reductase_MF521566.1”, which the MON15985 GMO was not known to contain.

**Figure 6.**
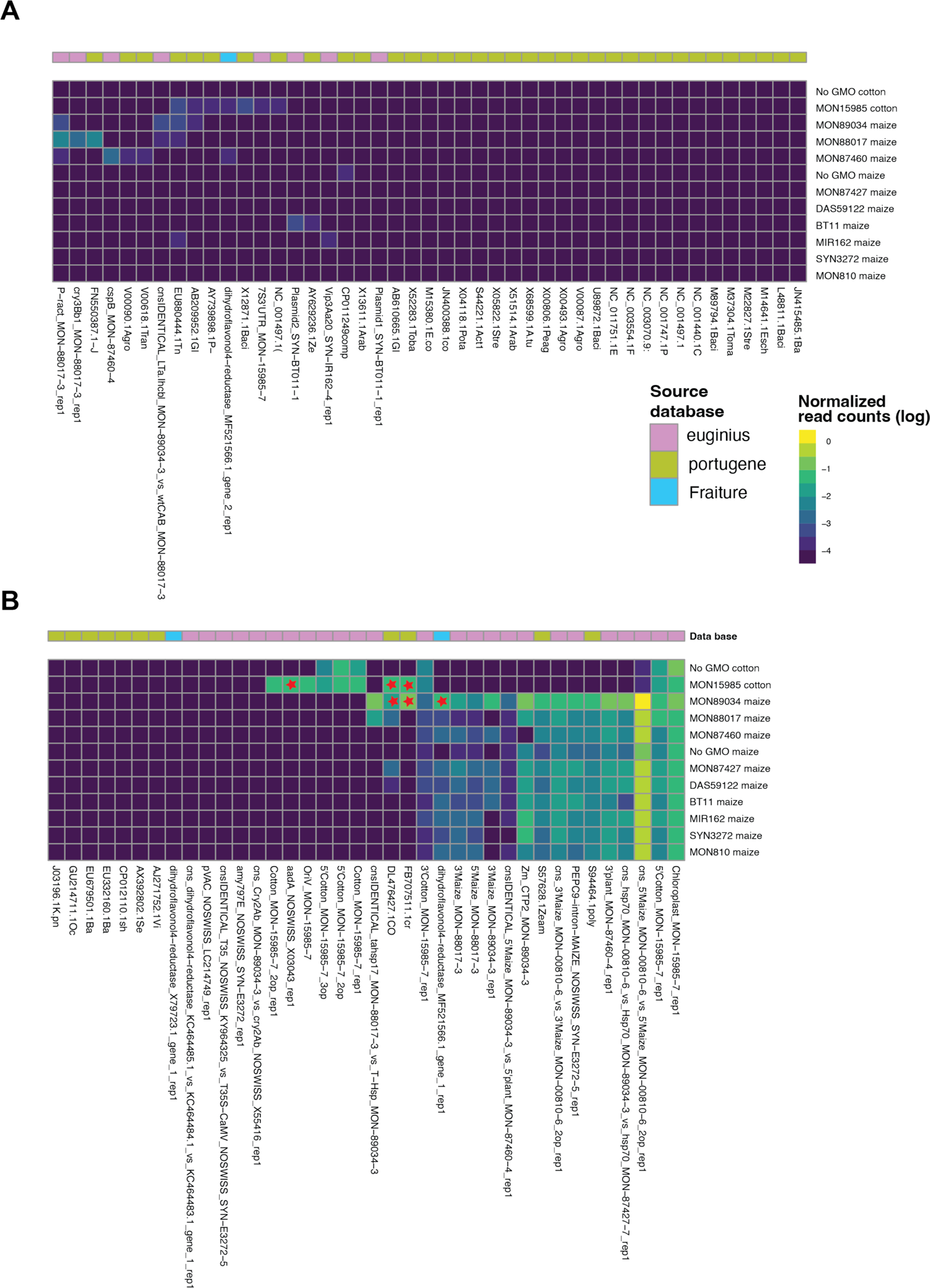
Amplifications of GM events are included in the databases euginius, portugene and Fraiture. Heatmap for 11 samples including nine different GMO events for maize with one non-GMO maize as control; one GMO event for cotton with one non-GMO cotton as control, aligned to the 82 different designed amplicons from the three databases (euginius, portugene and Fraiture) and read counts were normalized using barcoding loci sequencing depth. The red stars are marking amplification in the samples for MON15985 GMO cotton where “FB707511.1 cry1A.105”, “DL476427.1 CORN EVENT” and “aadA” were amplified; for the MON 89034 “FB707511.1 cry1A.105” and “DL476427.1 CORN EVENT” and “dihydroflavonol4-reductase_MF521566.1”.

## DISCUSSION

Quantative PCR has been for long the widely accepted standard for GMO detection in all matrices that could be circulating: food, feed, seeds and in the environment. qPCR is highly sensitivity and labour-intensive. Improving methods for detecting GMOs is therefore important. Here, we assessed the feasibility of a microfluidics-based approach for the detection of GM events. We desigend a set of 230 amplicons that represent 82 unique GM events. In addition, 27 plant barcoding primers allowed to determine species contained in the samples. The advantage of using amplicon sequencing is the versatility and expandability to perform monitoring of samples without prior knowledge of the genetic modifications present.

Primers to amplify the barcoding genes rbcL and ITS allowed to identify most plant species contained in the sample. Samples originating from otton were however not well identified withlow read counts mapping to the rbcL gene. Low coverage may be due to selected primers performing poorly in those samples. The complexity of amplifying a barcoding locus using multiple overlapping pairs of primers was apparent in the read mapping patterns along the rbcL and ITS sequences. Replacing the pool of barcoding loci primers with pairs of custom-designed primers for a specific range of species would alleviate the complexity in amplification and likely increase consistency in amplification across the desired species.

The detection of GM events in positive control samples was successful for a wide range of GM sequences. However, non-GMO controls amplified in some GM regions including 5’ and 3’ flanking sequences, the promoters, and terminators, which likely have homology to regions in the genome outside of the GM sequences. Such a lack of specificity in flanking and promoter sequences of GM events against regular plant DNA sequences can explain why GMO event promoters are also amplifying in non-GMO crops (Jores et al. 2021). The GM regions, which were amplified in non-GMO samples showed indeed sequence homology in genomes of non-GMO crops. This supports the lack of specificity in the 5’ and 3’ flanking sequences of GM events. However, the recovered homologous sequences in crop genomes did not present 100% identity, which suggests that SNP calling could be used to differentiate between reads mapping to a GM event versus reads mapping to non-GMO sequences elsewhere in the genome.

Previous studies describing multiplex detection of GM events were largely based on amplicons stemming from known primer pairs used in routine GMO food and feed sample screening strategies and approved in GMO reference material (Scholtens et al. 2017, Arulandhu et al. 2018). In some jurisdictions including the European Union, there is an obligation to proivde a detection method specific to the authorized GM event. As the common practice usually is centred on the use of qPCR, often only primer sequences are made available, limiting the investigation of GM events. The dependency on a certified set of primers restricts the completeness of the GM event sequences to be recovered, because amplicon sequences do not cover the complete GM event. The lack of long amplicon sequences and reference material accessibility are challenging to investigate GMO sequences comprehensively (Moreira, Carneiro, and Pereira 2017a). Our study shows that a *de novo* design of GM amplicons is feasible using public sequence information and that a broad range of potential GM sequences can be assessed in parallel. The approach of a microfluidics-based targeted amplicon sequencing assay enables to screen both hundreds of samples (or replicates) simultaneously but also allows for large sets of primers to be included in parallel. In principle, the microfluidics chips would allow the pooling of thousands of primer pairs for single-step amplifications. Given the uncertainty of amplifying specific sequences from unknown samples and modification events, the sequence information provides significantly greater certainty about the identity of an amplified sequence. This contrasts with qPCR approaches that lack direct validation capabilities under non-standard conditions. In contrast, targeted amplicon sequencing is less sensitive compared to qPCR. In our analyses, we found reliable amplification up to ∼1:100 dilutions, at lower concentrations the detection is likely to be obnly poorly reproducible. Such detection limitations can be remedied partially by increasing the overall sequencing coverage of the amplicons as sensitivity is at least partially correlated with sequencing depth.

This study presented a comprehensive amplicon sequencing assay that leverages the parallelization offered by microfluidics platforms and the depth of NGS for the detection of GMOs. Our work fits into recent efforts to standardize and propose a statistical framework for the detection of GMOs (Willems et al. 2016) based on the number of reads aligned per sample. Our workflow expands the capabilities by targeting a large number of sequences of interest specifically and allowing for the efficient detection of plant species present in a sample. With 230 designed amplicons corresponding to GM events and additional primers for species-specific barcoding, our approach represents a significant advancement in GMO detection. The assay robustness was demonstrated by successfully identifying ten known genetic modifications across different crop species and showcasing the potential for uncovering undocumented genetic events. The integration of rbcL and ITS barcoding primers enhanced the assay’s precision by enabling accurate species identification within mixed samples, an essential step in standardised GMO presence across samples. Challenges remain in distinguishing between GM events and homologous non-GMO sequences. The use of sequence information provided by NGS offers a direct validation of the detected genetic material. The successful identification of GMOs in this study underscores the importance of developing advanced screening methods. Our study shows the relevance of amplicon sequencing that can be realistically implemented into GMO detection and efficiently analyzed using a structured bioinformatics pipeline.

## Acknowledgments

Data was generated in collaboration with the Genetic Diversity Centre (GDC), ETH Zurich.

